# Dietary vitamin B12 deficiency impairs motor function and changes neuronal survival and choline metabolism after ischemic stroke in middle aged male and female mice

**DOI:** 10.1101/2021.08.17.456684

**Authors:** Gyllian B. Yahn, Brandi Wasek, Teodoro Bottiglieri, Olga Malysheva, Marie A. Caudill, Nafisa M. Jadavji

**Author notes:** **Corresponding Author:** Nafisa M. Jadavji, PhD, Department of Biomedical Sciences, College of Graduate Studies, College of Veterinary Medicine, College of Osteopathic Medicine, Midwestern University, 19555 N 59th Ave, Glendale, AZ 85308, USA.

## Abstract

Nutrition is a modifiable risk factor for ischemic stroke. As people age their ability to absorb some nutrients decreases, a primary example is vitamin B12. Older individuals with a vitamin B12 deficiency are at a higher risk for ischemic stroke and have worse stroke outcome. However, the mechanisms through which these occur remain unknown. The aim of the study was to investigate the role of vitamin B12 deficiency in ischemic stroke outcome and mechanistic changes in a mouse model. Ten-month-old male and female mice were put on control or vitamin B12 deficient diets for 4-weeks prior to and after ischemic stroke to the sensorimotor cortex. Motor function was measured, and tissues were collected to assess potential mechanisms. All deficient mice had increased levels of total homocysteine in plasma and liver tissues. After ischemic stroke, deficient mice had impaired motor function compared to control mice. There was no difference between groups in lesion volume, however, there was an increase in total apoptosis within the ischemic region of brain tissue in male deficient mice. More neuronal survival was present in ischemic brain tissue of the deficient mice compared to controls. Additionally, there were changes in choline metabolites in ischemic brain tissue because of a vitamin B12 deficiency. The data presented in this study confirms that a vitamin B12 deficiency worsens stroke outcome in male and female mice. The mechanisms driving this change may be a result of neuronal survival and compensation in choline metabolism within the damaged brain tissue.

## 1. Introduction

It is known that eating healthy is important for good health and can reduce the risk of diseases such as stroke [1]. Vitamin B-12, also referred to as cobalamin, is a water-soluble vitamin found naturally in foods such as red meat, poultry, eggs, dairy products, seafood, and fortified cereals [2]. It has very important roles in the human body as it is necessary for proper red blood cell formation, neurological function, myelin synthesis and repair, and DNA synthesis. A vitamin B-12 deficiency is a serious and common complication that is defined as low plasma and tissue levels of vitamin B-12 [3]. It can affect all age ranges, however it has a much higher prevalence within the elderly population [1], [4]. A deficiency is commonly caused by malabsorption, decreased acid secretion, and reduced intrinsic factor production [4]. Vitamin B-12 is an important co-factor in the conversion of homocysteine to methionine. Elevated levels of homocysteine are well associated with increased risk for cardiovascular disease, such as ischemic stroke [5].

In elderly patients, a vitamin B12 deficiency is commonly caused by malabsorption, decreased acid secretion, or reduced intrinsic factor production [4]. Having a deficiency in vitamin B12 results in a multitude of problems and can exacerbate outstanding conditions or developing conditions. The recommended daily amount of vitamin B12 is 2.4 µg and the average daily diet contains a range of 3-30 µg [6]. Interestingly, only 50% of patients who are vitamin B12 deficient present with low levels of serum vitamin B12, resulting in a substantial number of undiagnosed patients [2]. Clinical manifestations can be set back due to the high amounts of hepatic storage of vitamin B12 [6].

Adequate levels of vitamin B12 are needed for successful aging [7]. The presence of atrial fibrillation, vitamin B12 deficiency, and resultant elevated levels of plasma total homocysteine (tHcy) increases with age and are a risk factor for stroke [8], [9]. Clinical data strongly suggest that low levels of vitamin B12 are a risk factor for ischemic stroke and also impacts stroke outcome [10], [11]. There is a gap in the understanding of the mechanisms in which a vitamin deficiency leads to increased risk of stroke and worse stroke outcome. This study aims to assess the effects of a vitamin B12 deficiency on stroke outcome and related mechanisms.

## 2. Methods

### 2.1. Animals

All experiments in animals were approved by the Midwestern University IACUC committee. Female and male C57/BL6J mice were obtained for Jackson laboratories for this study. A total of 44 mice were obtained, 22 males and 22 females. The mice 10-months-old upon arrival and were habituated for 1 week prior to the start of experiments.

### 2.2. Experimental design

An overview of all experimental manipulations is outlined in Figure 1. After mice were habituated to the Midwestern University animal facility, they were randomly assigned to control or vitamin B12 deficient groups and placed on diets for 4 weeks. At 11 months of age, ischemic stroke was induced in animals using the photothrombosis model [12]–[16], the age of mice corresponds to middle aged in humans [17]. The ischemic stroke damaged the sensorimotor cortex, which allows for motor function assessment. Four weeks after damage motor function was measured in animals using the forepaw placement and accelerating rotarod tasks. After the completion of behavioral testing animals were euthanized and tissue was collected, including brain, liver, and blood tissue to study mechanisms. Brain tissue for cryosectioning was collected from animals, after they were perfused with phosphate buffered saline and 4% paraformaldehyde tissue. Cryosectioned brain tissue was used to quantify damage volume quantification, and immunofluorescence experiments. For one-carbon metabolite analysis in brain tissue, cortical ischemic damage and healthy tissue was micro dissected and snap frozen using dry ice. There were four experimental groups control male (MCon) and female (FCon); vitamin B-12 deficient male (MDef) and female (FDef).

**Figure 1.**
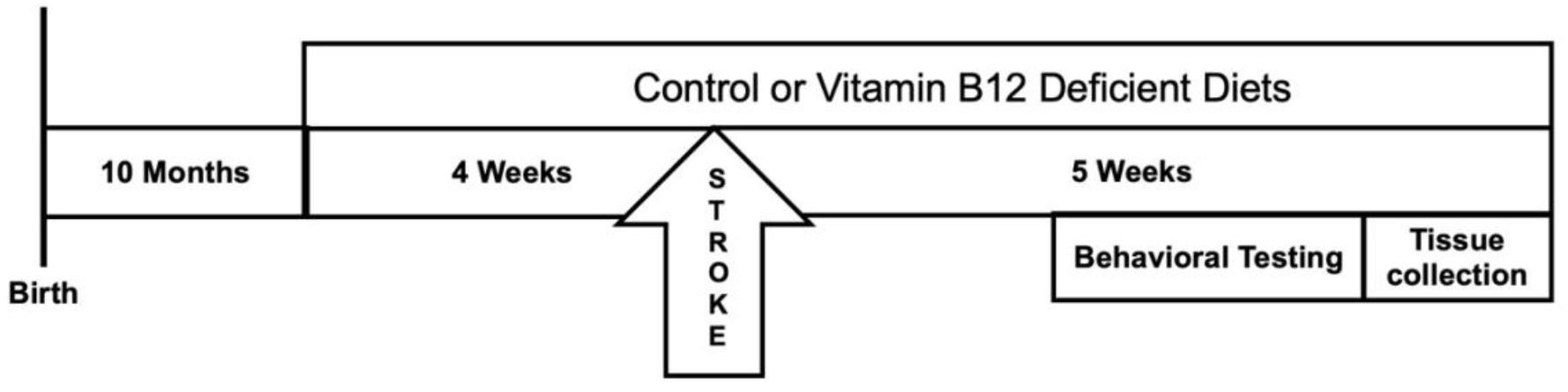
Timeline of experimental manipulation. Male and female C57Bl/6J mice arrived from Jackson Laboratories 10-months-old, and after one week of acclimation, animals were placed on a control or vitamin B-12 deficient diet for four weeks. Following the four weeks, ischemic stroke was induced using the Photothrombosis model in the sensorimotor cortex. After stroke animals were maintained on respective diets for 4 additional weeks, after which motor function of the mice was measured using the accelerating rotarod and forepaw placement tasks. At the completion of *in vivo* experiments animals were euthanized, and brain and liver tissue, as well as plasma was collected for further analysis.

### 2.3. Diet

The mice were placed on a vitamin B12 deficient (0 mg/kg) or a control (0.025 mg/kg vitamin B12) diet four weeks prior to photothrombosis damage and four weeks post photothrombosis damage. The diets were formulated by and purchased from Envigo. The control diet (TD. 190790) contains the recommended dietary amount of nutrients for mice [18]. The vitamin B12 deficient diet (TD. 190791) was formulated based on a previous study in mice that has shown it to be safe and have no negative side effects [19]. The mice had *ad libitum* access to food and water throughout the experiment. Body weights of each animal were recorded weekly.

### 2.4 Photothrombosis model

Using the photothrombosis model of ischemic stroke damage, mice were anesthetized with 4-5% isoflurane in oxygen. After anesthetization the mice had the top of their heads shaved and disinfected. Tear gel was used to prevent their eyes from drying out during while anesthetized.

0.03 mg/kg of Buprenorphine and one mL of saline were administered subcutaneously. Mice were then transferred to a stereotaxic apparatus (Harvard Apparatus) and maintained at 2-2.5% isoflurane. The mice were placed on a heating pad and a probe was rectally inserted to maintain a body temperature of 37°C. Prior to laser exposure, mice were intraperitoneally injected with 10 mg/kg of photoactive Rose Bengal (Sigma) followed by a 5-minute delay to allow the dye to enter circulation. Skin at the top of the head was surgically cut to expose the skull and then the sensorimotor cortex was targeted using stereotaxic coordinates (3 cm above, mediolateral + 0.24 mm from Bregma). The skull of the mice was exposed to a laser (Beta Electronics, wavelength: 532 nm) for 15 minutes. For recovery of post-operative pain, Buprenorphine was administered to all animals prior to ischemic damage.

### 2.5. Behavioral Testing

#### 2.5.1. Accelerating Rotarod

A standard accelerating rotarod apparatus (Harvard Apparatus) was used to measure walking movements and balance previously described [14], [20], [21]. Thirty centimeters above the ground, mice were placed on a rotating rod 3 cm in diameter and 6 cm wide in which the speed gradually increases from 4 to 60 revolutions per minute over 8 minutes. When mice fall off the rotarod, a digital sensor recorded the latency, in seconds, to fall off the accelerating rod. An average of three trials per mouse was taken with an inter trial interval of five minutes.

#### 2.5.2. Forepaw placement task

To measure spontaneous forepaw usage, mice were placed in a 19 cm high, 14 cm diameter cylinder, and the placement of their forepaws on the cylinder wall during natural exploratory rearing behaviors was recorded using a digital camera for frame-by-frame analysis [14], [22]. During a rear, the first forepaw placement on the wall was recorded as impaired, non-impaired, or both.

### 2.6. Total homocysteine levels

At the time of euthanization, blood was collected by cardiac puncture in EDTA coated tubes, centrifuged at 7000g for 7 minutes at 4C to obtain plasma. Liver tissue was also removed at the same time and samples were stored at -80C, until time of analysis. Total homocysteine (tHcy) in plasma and liver were measured by liquid chromatography tandem mass spectrometry (LC-MS/MS) as previously described ([23].

### 2.7. Brain tissue processing

Brain tissue was sectioned using a cryostat at 30 μm and slide mounted in serial order. There were six slides full of brain tissue sections of the damaged area per mouse and each animal had a minimum of four sections that were used for quantification. ImageJ (NIH) was used to quantify ischemic damage volume by measuring the area of damaged tissue [24].

### 2.8. Immunofluorescence experiments

Brain tissue was used in immunofluorescence analysis to assess molecular mechanisms and brain tissue staining was performed to investigate potential mechanisms. Primary antibodies included, active caspase-3 (1:100, Cell Signaling Technologies) to measure apoptosis and phospho-AKT (protein kinase B) (1:100, Cell Signaling Technologies). All brain sections were stained with a marker for neuronal nuclei, NeuN (1:200, AbCam). Primary antibodies were diluted in 0.5% Triton X and incubated with brain tissue overnight at 4°C. The next day, brain sections were incubated in Alexa Fluor 488 or 555 (Cell Signaling Technologies) and secondary antibodies were then incubated at room temperature for 2 hours and stained with 4’, 6-diamidino-2-phenylindole (DAPI) (1:1000, Thermo Fisher Scientific). The stains were analyzed using a microscope (Zeiss) and all images were collected at the magnification of 40X.

In brain tissue within the ischemic region, co-localization of active caspase-3 or phospho-AKT with NeuN labelled neurons were counted and averaged per animal. A positive cell was indicated by co-localization of the antibodies of interest located within a defined cell. Cells were distinguished from debris by identifying a clear cell shape and intact nuclei (indicated by DAPI and NeuN) under the microscope. All cell counts were conducted by two individuals blinded to treatment groups. The number of positive cells were counted in three brain sections per animal. For each section, three fields were analyzed. The number of positive cells were averaged for each animal.

### 2.9. Choline metabolites measurements

Frozen brain tissue from ischemic and non-ischemic cortex was measured for acetylcholine, betaine, choline, glycerophosphocholine, phosphocholine, phosphatidylcholine, and sphingomyelin levels using the LC-MS method as previously reported [25].

### 2.10. Data analysis and statistics

All data were analyzed by two individuals that were blinded to experimental treatment groups. GraphPad Prism 6.0 was used to analyze all data from the study. Two-way ANOVA analysis was performed when comparing the mean measurement of both sex and dietary group for behavioral testing, plasma tHcy measurements, lesion volume, immunofluorescence staining, and choline measurements. Significant main effects of two-way ANOVAs were followed up with Tukey’s post-hoc test to adjust for multiple comparisons. All data are presented as mean ± standard error of the mean (SEM). Statistical tests were performed using a significance level of 0.05.

## 3. Results

### 3.1 Behavioral Testing

#### 3.1.1. Accelerating Rotarod

After damaging the sensorimotor cortex, we measured the time it took for animals to fall of the accelerating rotarod (latency to fall). There was a sex main effect (Figure 2A, *p* = 0.02). Male mice were not able to stay on the ladder beam compared to female mice (Figure 6A, *p* = 0.03). There was no interaction between diet and sex (*p* = 0.90) and diet groups *(p* = 0.09).

**Figure 2.**
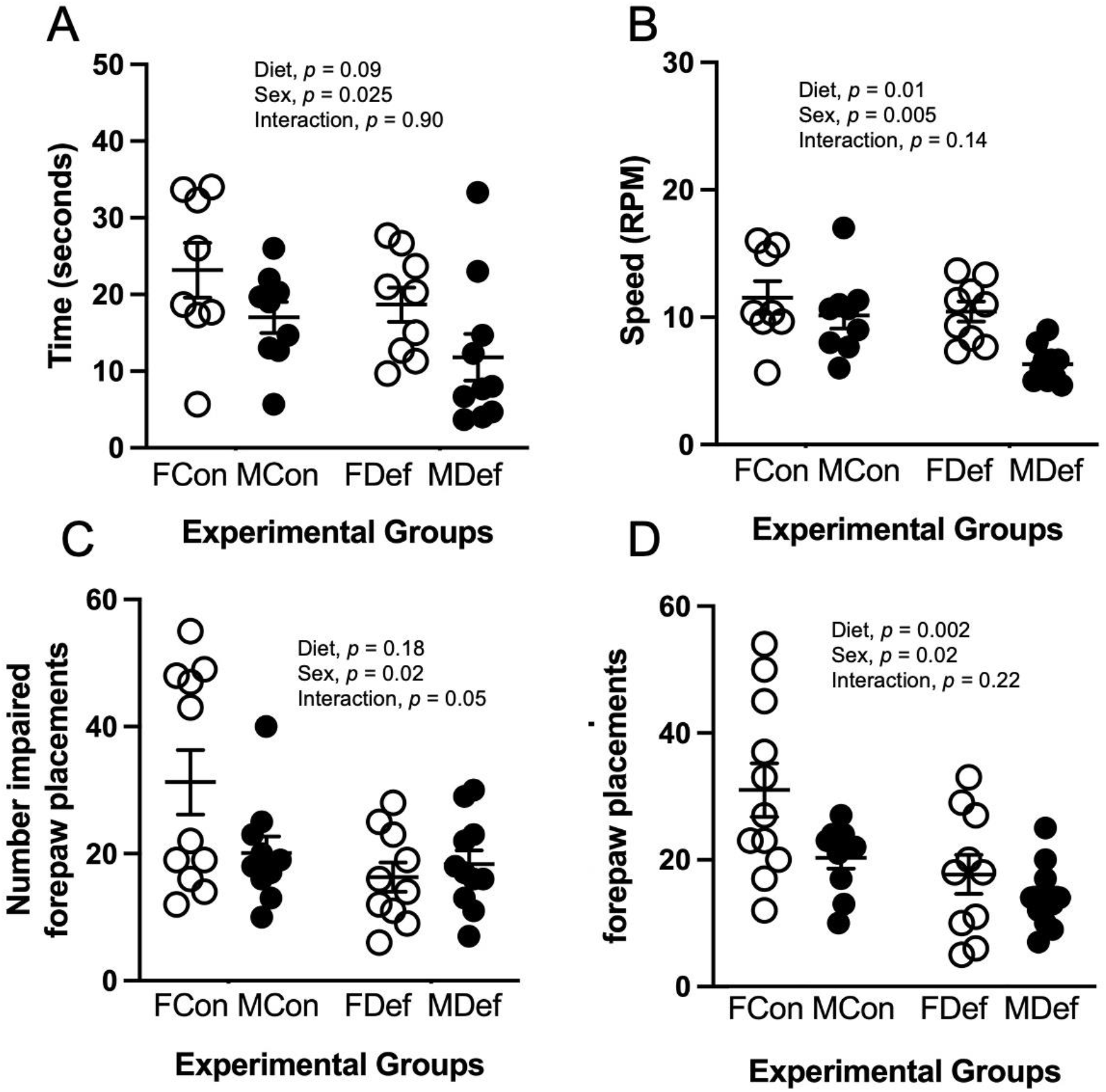
Motor function after ischemic stroke in male (M) and female (F) middle-aged mice fed B-12 deficient (Def) or control (Con) diet for 9 wk. Accelerating rotarod, the latency to fall (A) and revolutions per minute (B) were measured. The forepaw placement task number of impaired (A) and non-impaired (B) forepaw placements were recorded. Individual values, means, and SEMs are shown, n = 10 to 11.* *p* < 0.05 pairwise comparison between FCon and FDef mice.

In terms of speed achieved on the accelerating rotarod (revolutions per minute; RPM), there were differences in diet (Figure 2B, *p* = 0.01) and sex (*p* = 0.005). Control diet mice achieved higher levels of RPM compared to vitamin B-12 deficient mice (*p* = 0.014). MCon mice achieved a higher RPM compared to MDef animals (*p* = 0.02). FCon and MCon, as well as FDef mice achieved higher speeds compared to MDef mice (Figure 6B, *p* = 0.005). There were no interactions between diet and sex (*p* = 0.14).

#### 3.1.2. Forepaw Placement

The forepaw placement task evaluated forepaw usage through natural rearing behaviors of rodents [22]. Four weeks after ischemic damage to the sensorimotor cortex there was a difference in impaired forepaw usage scores between vitamin B-12 deficient diet animals and controls (Figure 2C, *p* = 0.02). FCon mice used their impaired forepaw more than vitamin B-12 deficient animals (*p* = 0.02). There was no difference between sex (*p* = 0.18) or interaction between sex and diet (*p* = 0.05).

Diet (Figure 2D, *p* = 0.002) and sex (*p* = 0.02) both affected non-impaired forepaw usage. FCon used their non-impaired forepaw more than FDef (*p* = 0.012) and MDef (*p* = 0.008). There was no interaction between diet and sex (*p* = 0.22) in non-impaired forepaw usage.

### 3.2 Total homocysteine levels

Vitamin B-12 deficiency resulted in an increase of plasma homocysteine levels (Figure 3A; *p* = 0.0006). FDef mice had elevated levels of homocysteine compared to FCon (*p* = 0.05). There was no sex difference (Figure 2A; *p* = 0.66) and no interaction between diet and sex for homocysteine concentrations (*p* = 0.76).

**Figure 3.**
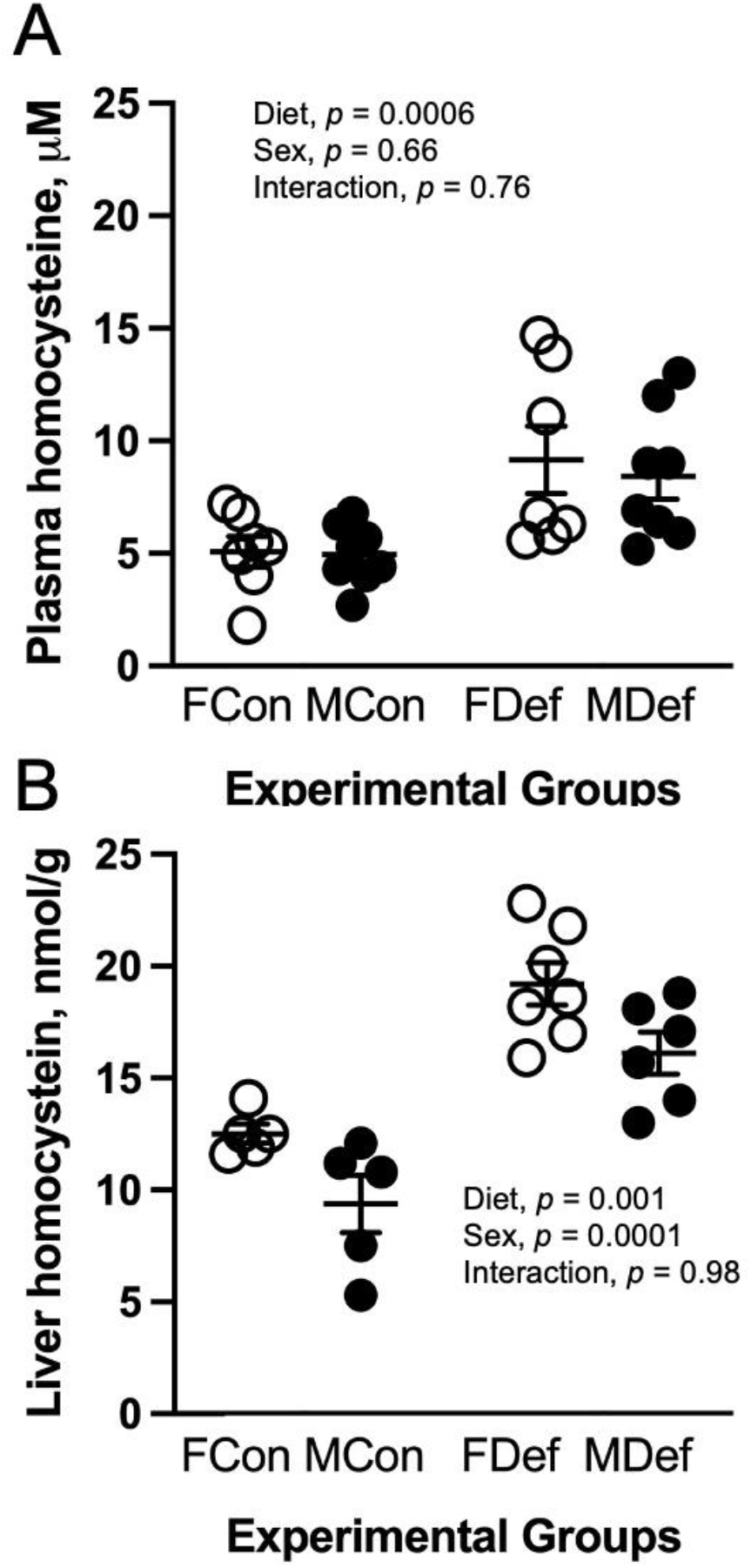
Plasma (A) and liver (B) total homocysteine concentrations in male (M) and female (F) middle-aged mice fed B-12 deficient (Def) or control (Con) diet for 9 wk. Individual values, means, and SEMs are shown, n = 7 - 8. *** *p* < 0.01 pairwise comparison between FCon and FDef or MCon and MDef mice.

In vitamin B-12 deficient mice, liver tHcy levels were elevated compared to control diet mice (Figure 3B, *p* = 0.0001) and there was also a sex effect (Figure 2B, *p* = 0.005). FDef mice had elevated levels of tHcy compared to FCon (*p* = 0.001). In liver tissue, there was no interaction between diet and sex in homocysteine levels (*p* = 0.98).

### 3.3 Ischemic Damage Volume Quantification

Representative images of ischemic damage to the sensorimotor cortex are shown in Figure 4A. Quantification of damage volume revealed that there was no interaction between sex and diet (Figure 4B, *p* = 0.08) and diet (*p* = 0.08) or sex (*p* = 0.28) main effects.

**Figure 4.**
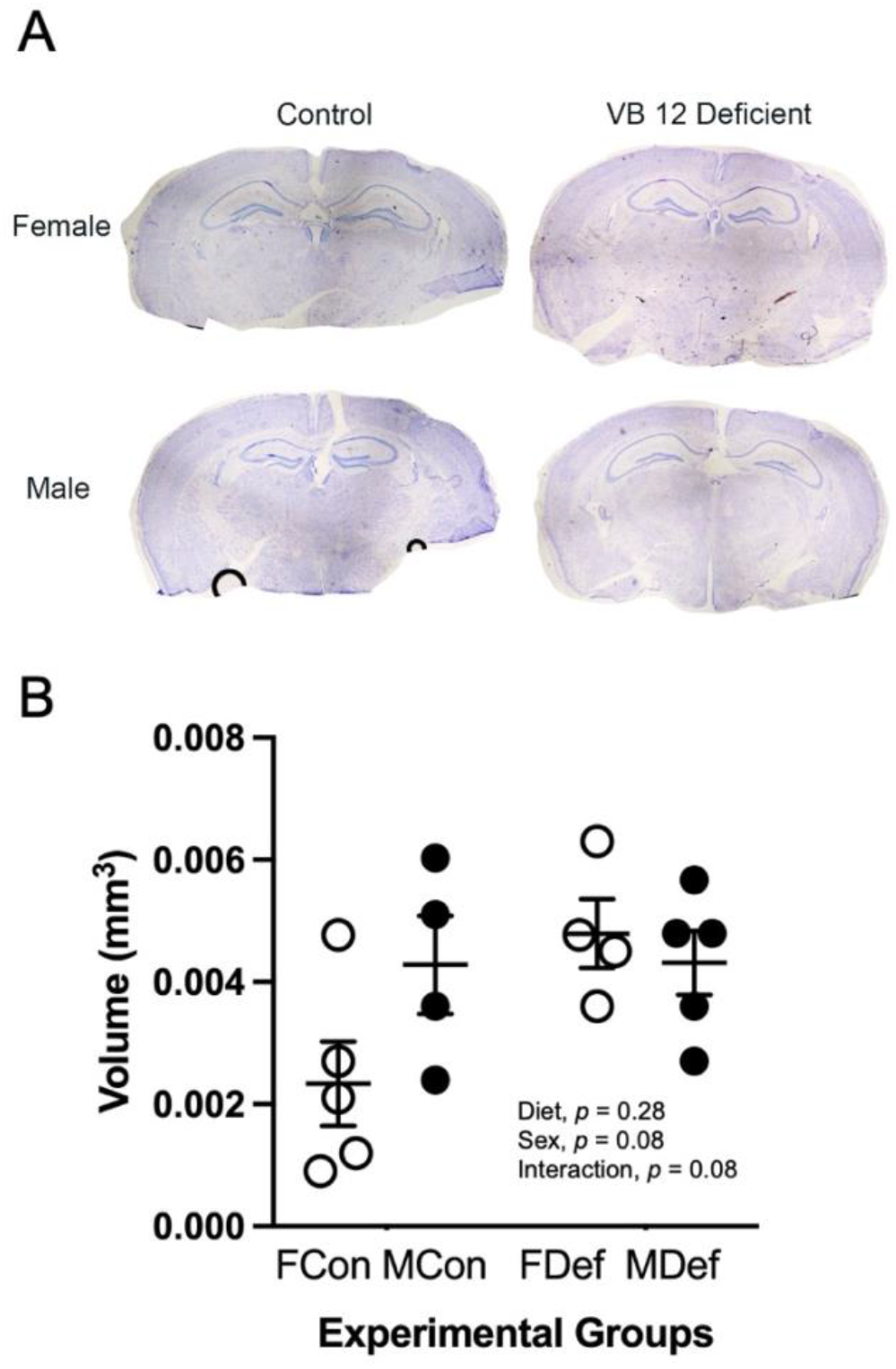
Ischemic damage volume in male (M) and female (F) middle-aged mice fed B-12 deficient (Def) or control (Con) diet for 9 wk. Representative cresyl violet image (A) and ischemic damage volume quantification (B). Individual values, means, and SEMs are shown, n = 4 to 5.

### 3.4 Brain Tissue Immunofluorescence Staining

#### 3.4.1. Apoptosis

Representative images of neuronal active caspase-3 staining from all groups are shown in Figure 5A. Quantification of neuronal apoptosis revealed that there was no interaction (Figure 5B, *p* = 0.17) and diet (*p* = 0.06) or sex (*p* = 0.12) main effects.

**Figure 5.**
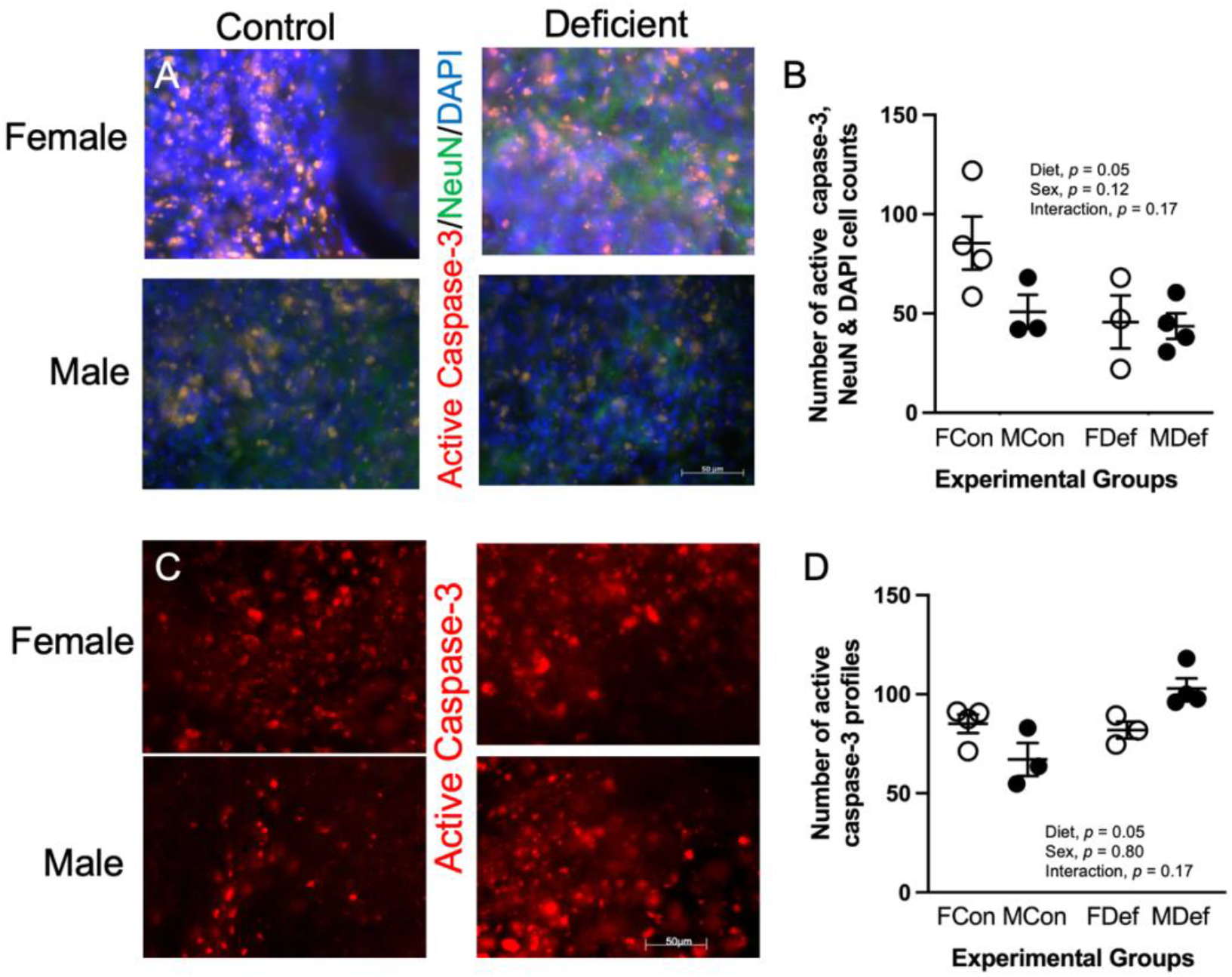
Active caspase-3 expression within ischemic region in male (M) and female (F) middle-aged mice fed B-12 (Def) or control (Con) diet for 9 wk. Representative images of immunofluorescence staining with positive semi quantitative spatial co-localization of active caspase-3 with neuronal nuclei (NeuN) and 4′,6-diamidino-2-phenylindole (DAPI) (A). Quantification of active capsapse-3, NeuN, and DAPI cell counts (B). Individual values, means, and SEMs are shown, n = 3 to 4. The scale bar = 50 μm. *** *p* < 0.01 pairwise comparison between MCon and MDef mice.

Representative images of total active caspase-3 within the damage area are shown in Figure 5C. There was a difference between control and vitamin B-12 deficient animals (Figure 5D, *p* = 0.02). MDef animals had more total active caspase-3 cells within the damage area compared to MCon mice (*p* = 0.006). There was no interaction (*p* = 0.17) and sex main effect (*p* = 0.80).

##### 3.4.2.1. Neuronal Cell Survial and Proliferation

Representative images of neuronal survival and proliferation from all groups are shown in Figure 6A. There was a difference between vitamin B-12 deficient and control diet animals (Figure 6B; *p* = 0.003), MDef animals had more positive neuronal pAKT (*p* = 0.05). There was no interaction (*p* = 0.27) or sex effect (*p* = 0.86).

**Figure 6.**
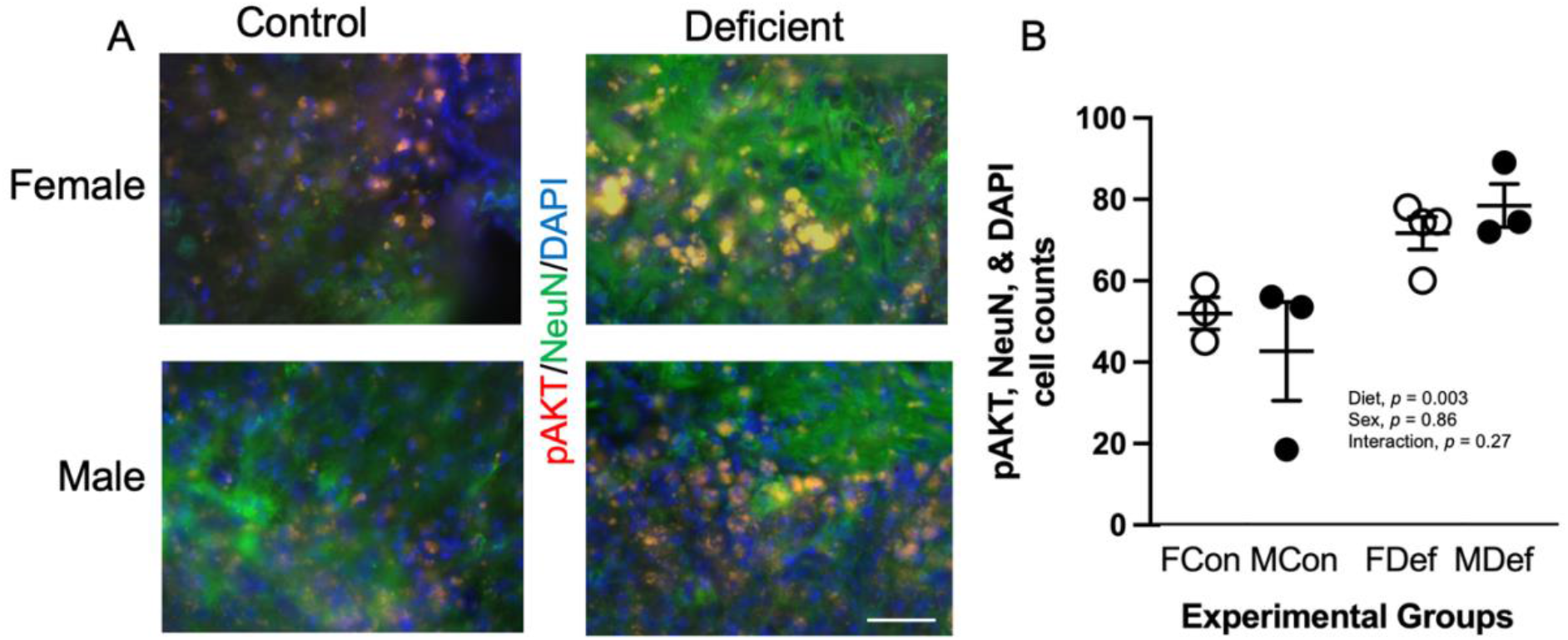
Phospho-AKT (pAKT) expression within ischemic region in male (M) and female (F) middle-aged mice fed B-12 (Def) or control (Con) diet for 9 wk. Representative images of immunofluorescence staining with positive semi quantitative spatial co-localization of active caspase-3 with neuronal nuclei (NeuN) and 4′,6-diamidino-2-phenylindole (DAPI) (A). Quantification of pAKT, NeuN, and DAPI cell counts (B). Individual values, means, and SEMs are shown, n = 3 to 4. The scale bar = 50 μm. * *p* < 0.05 pairwise comparison between MCon and MDef mice.

### 3.5. Cortical Brian Tissue One-Carbon Metabolite Measurements

Methionine levels were decreased in vitamin B-12 deficient brain tissue compared to controls (Table 1, diet effect, *p* = 0.02). Betaine levels were reduced in MDef and MCon diet animals compared to females (sex main effect, *p* = 0.003; diet main effect, *p* = 0.04). In ischemic damaged brain tissue, there was no interactions in any brain tissue between diet and sex for any of the one carbon metabolites measured.

**Table 1.** One-carbon metabolites concentration in cortical ischemic brain tissue of male and female mice maintained on control or vitamin B-12 deficient diets.^1^

| Metabolites<br>($\mu\text{mol/g}$) | FCon | MCon | FDef | MDef | Diet x<br>Sex | Diet | Sex |
| --- | --- | --- | --- | --- | --- | --- | --- |
| Methionine | $0.06 \pm 0.13$ | $0.09 \pm 0.06$ | $0.01 \pm 0.02$ | $0.01 \pm 0.12$ | $p = 0.40$ | $p = 0.02$ | $p = 0.15$ |
| Acetylcholine | $0.004 \pm 0.009$ | $0.005 \pm 0.0005$ | $0.006 \pm 0.00002$ | $0.004 \pm 0.003$ | $p = 0.18$ | $p = 0.70$ | $p = 0.73$ |
| Betaine | $0.02 \pm 0.001$ | $0.01 \pm 0.0009$ | $0.02 \pm 0.002$ | $0.01 \pm 0.002$ | $p = 0.21$ | $p = 0.04$ | $p = 0.003$ |
| Choline | $0.07 \pm 0.004$ | $0.05 \pm 0.002$ | $0.06 \pm 0.007$ | $0.04 \pm 0.007$ | $p = 0.06$ | $p = 0.94$ | $p = 0.50$ |
| GPC | $0.63 \pm 0.019$ | $0.76 \pm 0.060$ | $0.68 \pm 0.03$ | $0.77 \pm 0.06$ | $p = 0.90$ | $p = 0.93$ | $p = 0.08$ |
| Phosphocholine | $0.38 \pm 0.03$ | $0.33 \pm 0.03$ | $0.34 \pm 0.02$ | $0.33 \pm 0.01$ | $p = 0.37$ | $p = 0.28$ | $p = 0.20$ |
| Phosphatidylcholine | $34.2 \pm 0.57$ | $35.6 \pm 0.57$ | $34.7 \pm 0.65$ | $35.6 \pm 0.23$ | $p = 0.57$ | $p = 0.59$ | $p = 0.034$ |
| Sphingomyelin | $4.27 \pm 0.19$ | $4.08 \pm 0.20$ | $4.27 \pm 0.07$ | $4.00 \pm 0.02$ | $p = 0.80$ | $p = 0.80$ | $p = 0.14$ |

<sup>1</sup>Values are means ± SEMs of 5 animals per group. Abbreviations: FCon, female control; FDef, female deficient; GPC, glycerophosphocholine; MCon, male control, MDef, male deficient.

In the non-ischemic cortical brain tissue, there was no interactions in any brain tissue between diet and sex for any of the one metabolite measured (Table 2). Acetylcholine levels were increased in vitamin B-12 compared control diet animals (diet main effect, *p* = 0.03).

**Table 2.**
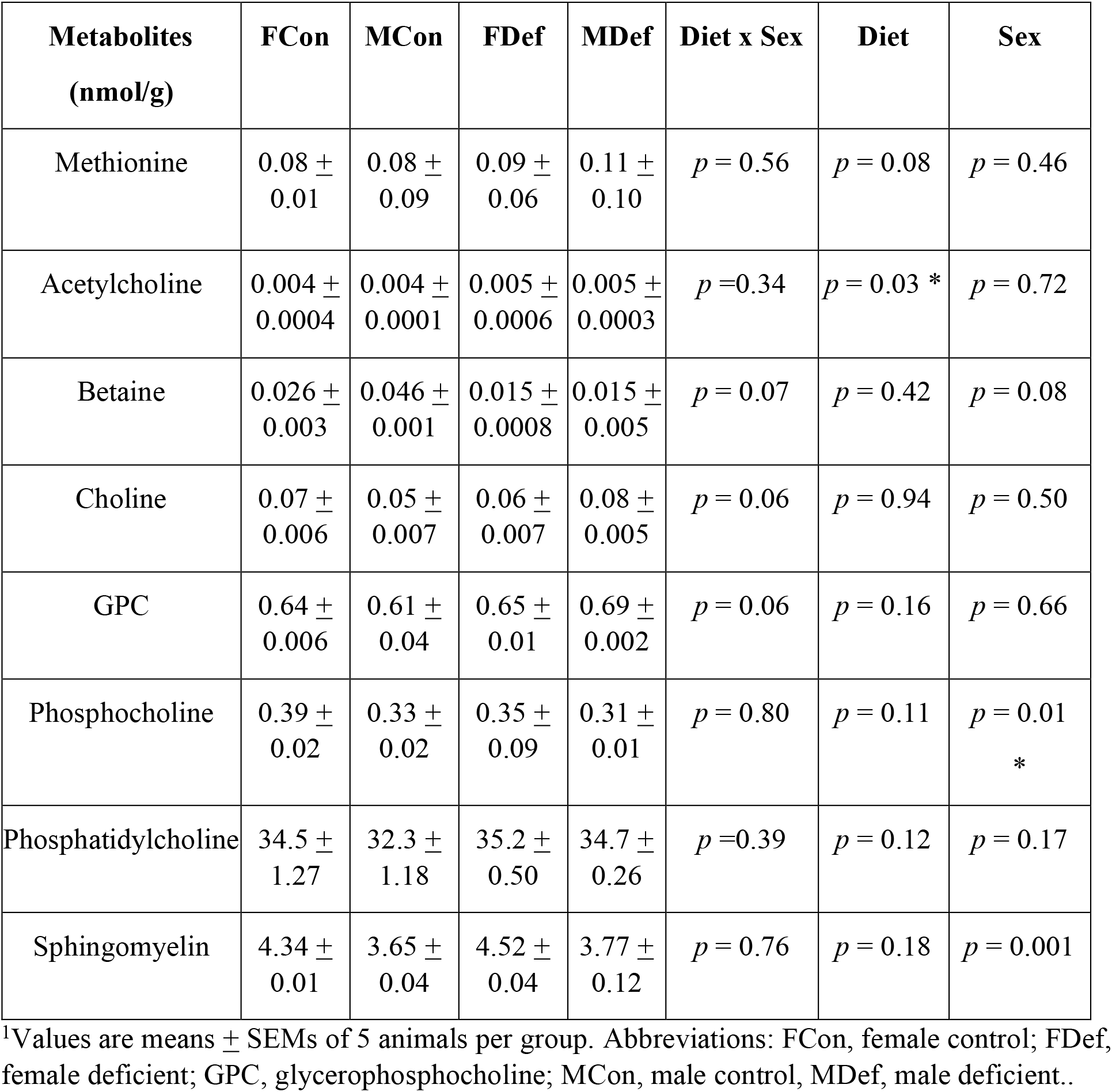
One-carbon metabolites concentration in cortical non-ischemic brain tissue of male and female mice maintained on control and vitamin B-12 diets.^1^

| **Metabolites**<br>**(nmol/g)** | **FCon** | **MCon** | **FDef** | **MDef** | **Diet x Sex** | **Diet** | **Sex** |
| --- | --- | --- | --- | --- | --- | --- | --- |
| Methionine | 0.08 ± 0.01 | 0.08 ± 0.09 | 0.09 ± 0.06 | 0.11 ± 0.10 | *p* = 0.56 | *p* = 0.08 | *p* = 0.46 |
| Acetylcholine | 0.004 ± 0.0004 | 0.004 ± 0.0001 | 0.005 ± 0.0006 | 0.005 ± 0.0003 | *p* = 0.34 | *p* = 0.03 * | *p* = 0.72 |
| Betaine | 0.026 ± 0.003 | 0.046 ± 0.001 | 0.015 ± 0.0008 | 0.015 ± 0.005 | *p* = 0.07 | *p* = 0.42 | *p* = 0.08 |
| Choline | 0.07 ± 0.006 | 0.05 ± 0.007 | 0.06 ± 0.007 | 0.08 ± 0.005 | *p* = 0.06 | *p* = 0.94 | *p* = 0.50 |
| GPC | 0.64 ± 0.006 | 0.61 ± 0.04 | 0.65 ± 0.01 | 0.69 ± 0.002 | *p* = 0.06 | *p* = 0.16 | *p* = 0.66 |
| Phosphocholine | 0.39 ± 0.02 | 0.33 ± 0.02 | 0.35 ± 0.09 | 0.31 ± 0.01 | *p* = 0.80 | *p* = 0.11 | *p* = 0.01<br>* |
| Phosphatidylcholine | 34.5 ± 1.27 | 32.3 ± 1.18 | 35.2 ± 0.50 | 34.7 ± 0.26 | *p* = 0.39 | *p* = 0.12 | *p* = 0.17 |
| Sphingomyelin | 4.34 ± 0.01 | 3.65 ± 0.04 | 4.52 ± 0.04 | 3.77 ± 0.12 | *p* = 0.76 | *p* = 0.18 | *p* = 0.001 |

<sup>1</sup>Values are means ± SEMs of 5 animals per group. Abbreviations: FCon, female control; FDef, female deficient; GPC, glycerophosphocholine; MCon, male control, MDef, male deficient..

Phosphocholine (sex main effect, *p* = 0.01) and sphingomyelin (sex main effect, *p* = 0.001) levels were decreased in males compared to female animals.

## 4. Discussion

Nutrition is a modifiable risk factor for stroke [28]. The elderly population is at higher risk of stoke as well as a vitamin B12 deficiency [4]. Worse stroke outcome has been reported in patients with a vitamin B12 deficiency [10], [11]. The aim of the study was to investigate the role of vitamin B12 deficiency in ischemic stroke outcome and mechanistic changes in a mouse model. Our findings showed that vitamin B12 deficient mice had more impairments in coordination and balance compared to control diet mice after ischemic damage. Forepaw usage was decreased in female mice maintained on the vitamin B12 diet. Male mice on a vitamin B12 deficient diet showed impaired coordination and balance. In both plasma and liver there was an increase in plasma and liver tHcy because of dietary vitamin B12 deficiency. There was no difference in the damage size and neuronal apoptosis induced by the ischemic stroke between groups. There was an increase in overall apoptosis within the ischemic cortex. We also report increased neuronal survival in vitamin B12 deficient male and female mice, along with changes in choline metabolism in brain tissue.

Lacunar stroke patients with vitamin B12 deficiency reported significantly worse stroke outcome, including fatigue and depressive-like symptoms [29]. A common stroke outcome is impaired motor function, and the two most common motor deficits from stroke are spasticity and paresis, resulting in decreased usage of that limb. We measured forepaw usage and saw that female mice had significantly reduced impaired and non-impaired forepaw usage. Male vitamin B12 deficient mice showed impairments in coordination and balance using the rotarod task. Our preclinical study results are consistent with clinical observations in terms of worse outcome after ischemic stroke when patients are vitamin B12 deficient [11].

In the present study, we did not observe any differences in damage volume between dietary groups; however, this is a gross measurement. In our immunofluorescence experiments we report a difference in in apoptosis between dietary groups. In our neuronal apoptosis data, there was a decrease in neuronal death in vitamin B12 deficient mice. This may be due to increased neuronal plasticity in the damaged area [30]. However, total levels of apoptosis showed an increase in overall positive active caspase-3 levels in male vitamin B12 deficient mice. Increased levels of apoptosis have been reported in clinical studies, for example, in an autopsy cohort consisting of 13 cases of fatal ischemic stroke, TUNEL-labelled cells with apoptotic morphology were disproportionately more frequent in the ischemic core, compared percentage of the cells in the non-damaged hemisphere [31]. Furthermore, in post-ischemic stroke there is elevated levels of neuronal death due to apoptosis in the damaged area of the brain [32]. A potential reason there is not an increase in apoptosis in female mice is the neuroprotective effects of estrogen and progesterone [33]. This finding could also reflect an increased vulnerability of glial cells, the other type of cell in the brain. Overall, these findings support clinical data being reported [11].

One-carbon metabolism incorporates several vitamins and nutrients. Other studies have shown that vitamin B12 status affects choline status in animals [26], [27]. We measured levels of choline in brain tissue in ischemic and non-ischemic cortical tissue. Our results show that ischemic brain tissue has higher methionine levels in vitamin B12 deficient mice. The increase in methionine levels might be in compensation to the dietary vitamin B12 deficiency, since the brain is sensitive to methionine [34]. We also report lower levels of betaine in ischemic brain tissue of vitamin B12 deficient mice, which might reflect what’s happening in liver and plasma, and availability for uptake by brain.

Part of this study investigated sex differences in response to ischemic stroke between dietary groups. This is an important component of preclinical studies as there have been a low number of studies using female mice [35], [36]. In our study, male and female mice maintained on vitamin B12 deficiency had impaired stroke outcome, although they had impairments of different motor tasks. In tissue, male mice showed more total apoptosis, however in our present study our data shows both males and females had increased neuronal survival within damage region in brain tissue. This is an interesting finding considering research has shown that men are more susceptible to a vitamin B12 deficiency [37]. The results of our study emphasizes the importance of studying of sex differences in preclinical stroke experiments and the possibility of individualized medicine [38].

## Acknowledgments

The authors would like to acknowledge the Chelsea Adamson for her assistance in photothrombosis surgeries.

## Data Availability Statement

The data that supports the findings of this study are available in the supplementary material of this article.

## Notes on Contributors

Gyllian B. Yahn MBS, is a Doctor of Dental Medicine student at Midwestern University. Gyllian graduated undergrad with a degree in Biology and Neuroscience and completed her Masters of Biomedical Science

Brandi Wasek is a Research Assistant at Baylor Scott & White Health

Teodoro Bottiglieri, PhD is the Program Director at the Center of Metabolomics at Baylor Scott and White Health.

Olga Malysheva, MSc is a Research Support Specialist in the Diviion of Nutritional Sciences at Cornell University.

Marie A. Caudill, PhD is a Professor in Diviion of Nutritional Sciences at Cornell University.

Nafisa M. Jadavji PhD is an Assistant Professor in Biomedical Sciences at Midwestern University (US) and Research Assistant Professor in Neuroscience at Carleton University (Canada).

